# Overcoming the preferred orientation problem in cryoEM with self-supervised deep-learning

**DOI:** 10.1101/2024.04.11.588921

**Authors:** Yun-Tao Liu, Hongcheng Fan, Jason J. Hu, Z. Hong Zhou

**Affiliations:** Department of Microbiology, Immunology, and Molecular Genetics, University of California, Los Angeles, CA, USA; California NanoSystems Institute, University of California, Los Angeles, CA, USA; Current address: Department of Molecular and Cell Biology, University of California, Berkeley, CA, USA

## Abstract

While advances in single-particle cryoEM have enabled the structural determination of macromolecular complexes at atomic resolution, particle orientation bias (the so-called “preferred” orientation problem) remains a complication for most specimens. Existing solutions have relied on biochemical and physical strategies applied to the specimen and are often complex and challenging. Here, we develop spIsoNet, an end-to-end self-supervised deep-learning-based software to address the preferred orientation problem. Using preferred-orientation views to recover molecular information in under-sampled views, spIsoNet improves both angular isotropy and particle alignment accuracy during 3D reconstruction. We demonstrate spIsoNet’s capability of generating near-isotropic reconstructions from representative biological systems with limited views, including ribosomes, β-galactosidases, and a previously intractable hemagglutinin trimer dataset. spIsoNet can also be generalized to improve map isotropy and particle alignment of preferentially oriented molecules in subtomogram averaging. Therefore, without additional specimen-preparation procedures, spIsoNet provides a general computational solution to the preferred orientation problem.

## Main

Recent advances in cryogenic electron microscopy (cryoEM) hardwares^1–4^ and imaging processing algorithms^5–8^ have enabled atomic structure determination of well-behaved macromolecular complexes in their near-native states^3, 4^, transforming cryoEM as a mainstream technique of structural biology. In the ideal situation, biological complexes are evenly distributed with random orientations within a thin layer of vitreous ice on a cryoEM grid^9^. However, such an ideal situation is rarely the case; instead cryoEM specimens typically suffer from the so-called “preferred” orientation problem, which is characterized by uneven or biased distribution of particle orientations^10^. It is now recognized that preferred orientation generally arises from the interaction of macromolecules with the air-water interface (AWI) or the support film-water interface during specimen preparation^11–13^ (Fig. 1a). Because the electrostatic and hydrophobicity properties vary across the surface of the protein, certain protein surfaces are preferred to be exposed on the AWI^14–16^. This preferred binding to the AWI leads to the orientation bias^15^.

**Figure. 1.**
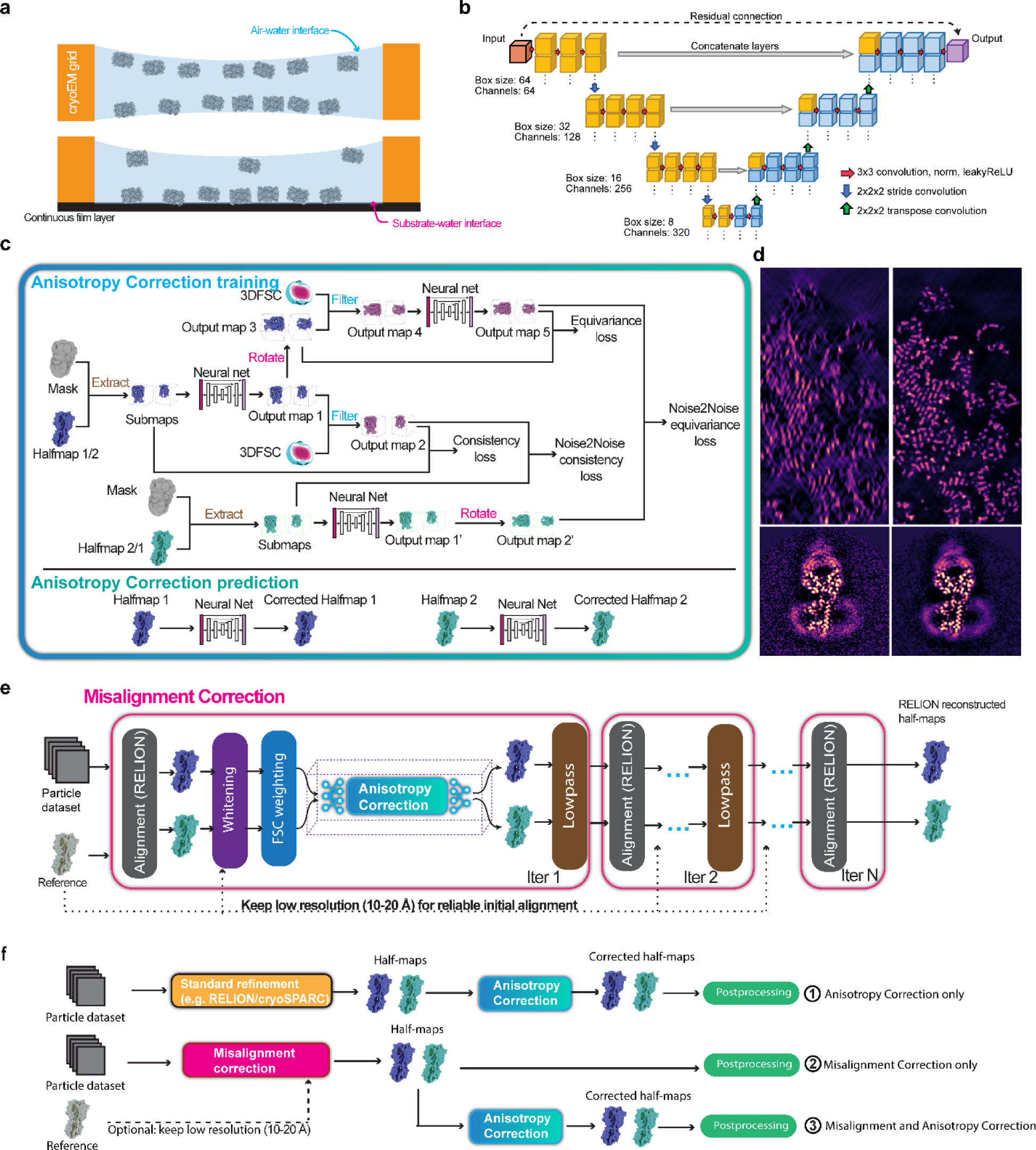
| The workflow of spIsoNet. **a,** Illustrations of two common scenarios that particles exhibit preferred orientations on cryoEM grids. **b,** Architecture of the U-net neural network used in spIsoNet. **c,** A diagram showing *Anisotropy Correction* algorithm highlighting the four-term loss functions. **d,** 2D slices of maps before and after spIsoNet *Anisotropy Correction*. Top panel shows a subregion of simulated ribosome. Bottom panel shows the denoising effect of spIsoNet with EMD-20806^55^. **e,** A diagram showing *Misalignment Correction*. **f,** Three workflows for using spIsoNet to tackle the preferred orientation problem.

The preferred orientation problem can severely compromise the quality and accuracy of the three-dimensional (3D) reconstructions, especially for low-symmetry or asymmetric macromolecules, resulting in artifacts in the reconstruction, such as skewed secondary structures, broken peptide or nucleotide chains, and distorted sidechain or nucleobase densities^17^. In some severe cases, the preferred orientation problem leads to a reconstructed map with distorted or incorrect shape^10^, which deteriorates the alignment of particles during the 3D refinement process, resulting in distortions/artifacts and even failure of obtaining a reconstruction. These artifacts misrepresent the true structure of the macromolecules and make atomic modeling difficult. Therefore, overcoming the prevailing challenge of preferred orientation is crucial for achieving high-throughput and accurate cryoEM structure determination, which will expand the applicability of cryoEM to determine structures of difficult biological specimens, understand molecular processes, and accelerate efforts for efficient structure-guided drug design.

Past efforts to address the preferred orientation problem have resorted to biochemical or physical approaches. Biochemical solutions include occupying the air-water interface with detergents^12, 14, 18^ or chemically modifying the protein specimen^19^, but are specimen-specific, typically require exhaustive screening, and may be detrimental to specimen quality. Physical approaches to alleviating the preferred orientation problem abound, including coating cryoEM grids with support films^11, 20–24^, using time-resolved vitrification devices^25–29^, collecting data from thicker ice regions^30^, and tilting the grids during image acquisition^10, 31^. However, these physical methods may be labor-intensive, expensive, and often have unintended consequences, such as higher background noise in images and a different type of preferred orientation induced by a support film-water interface. Therefore, the preferred orientation problem lacks a general and easy solution due to the specificity and complexity of existing methods.

Here, we introduce a self-supervised deep-learning method called single-particle IsoNet (spIsoNet) to restore angular isotropy in specimens that exhibit preferred orientations. spIsoNet addresses both anisotropic reconstruction and particle misalignment caused by the preferred orientation problem. spIsoNet consists of two modules, (1) map *Anisotropy Correction* and (2) particle *Misalignment Correction*. When applied to single-particle cryoEM, spIsoNet generated better reconstructions for various biological macromolecules—such as a 3.5 Å reconstruction from a non-tilted hemagglutinin dataset, which was previously intractable—by improving alignment accuracy and angular isotropy. When applied to subtomogram averaging of uneven orientation distribution, it also improves quality of the reconstructed map. The spIsoNet computational pipeline is specimen-independent and offers a much sought-after solution to the preferred orientation problem in single-particle cryoEM and single-particle cryogenic electron tomography (i.e., subtomogram averaging).

## Results

### Strategy of spIsoNet to address the preferred orientation problem

Even in anisotropic reconstructions, maps generated by cryoEM contain rich and recurring information from structural features such as secondary structures, amino acids, and atoms in well-sampled views. The organization of these features is constrained by hydrogen bonds, van der Waals forces, electrostatic and other interactions to minimize free energy, in a manner consistent with biophysical laws. With spIsoNet, we demonstrate that it is possible with modern deep learning methods to recover under-sampled information by merging information from well-sampled views of similar features present in the reconstruction, thus remedying the preferred orientation problem.

The preferred orientation problem has two major ramifications that hinder single-particle analysis. The first ramification is anisotropic reconstruction. By the central slice theorem, the Fourier transform of a 2D projection (i.e., a particle image) equals a 2D slice in 3D Fourier transform of the cryoEM map at its corresponding orientation^32^. A uniformly distributed particle orientation corresponds to a well-sampled 3D Fourier space, whereas an uneven particle orientation distribution corresponds to uneven sampling in Fourier space, causing a distorted 3D reconstruction in real space. The second ramification is particle misalignment. Standard cryoEM workflows require iterative particle orientation determination or alignment, where the maps reconstructed from one iteration are used as references for the next iteration^5, 7^. Thus, distorted maps reconstructed from the particles suffering from the preferred orientation problem lead to errors in particle alignment in the following iterations.

In spIsoNet, we implemented two modules—(i) *Anisotropy Correction* (Fig. 1c, d) and (ii) *Misalignment Correction* (Fig. 1e)—to remedy these ramifications of the preferred orientation problem, respectively. These modules can be used alone as a single command, or in tandem—first misalignment correction and then anisotropy correction to overcome the preferred orientation problem as demonstrated below (Fig. 1f).

### The Anisotropy Correction module

The *Anisotropy Correction* module requires two half-maps, a 3D Fourier shell correlation (3DFSC) volume^10^, and solvent mask as input. For user convenience, we implement a fast 3DFSC algorithm in spIsoNet. Regions in a 3DFSC volume with low intensity values indicate that these regions in the Fourier space of a map are under-sampled, while an isotropic map will generate an isotropic 3DFSC volume. Hence, we use 3DFSC as a proxy of directional resolution and density isotropy. Applying an anisotropic 3DFSC volume as a filter on a map in Fourier space approximates a cryoEM map with uneven orientation sampling.

The *Anisotropy Correction* module trains a deep neural network of the U-net architecture^33^ (Fig. 1b). The inputs of this module are the two half-maps from a standard cryoEM reconstruction or subtomogram averaging pipeline (Fig. 1c). Each half-map is randomly divided into smaller submaps as training data of spIsoNet. The loss function of the network training is defined as the weighted sum of four losses: consistency loss^34^, equivariance loss^34^, noise2noise^35^ consistency loss, and noise2noise equivariance loss. By minimizing this loss function, the neural network simultaneously accomplishes denoising and information recovery (Fig. 1d, Extended Data Fig. 1). The conceptual design of consistency and equivariance losses is based on the end-to-end, self-supervised framework of equivariant imaging proposed previously^34^.

Minimizing consistency loss ensures the 3DFSC-filtered, network-predicted map approximates the original map; therefore, the recovered information will only reside in the original under-sampled area in the Fourier space. To calculate the consistency loss, each extracted submap is passed through the neural network to generate the initial output map (referred to as output map 1). Output map 1 is then filtered by the 3DFSC filter, resulting in a 3DFSC-filtered and network-predicted map (output map 2). Consistency loss is defined as the difference between the output map 2 and the input submap.

By minimizing the equivariance loss, the network learns the how to recover information leveraging the fact that characteristic features, such as amino acids and nucleic acids, should be similar regardless of their positions and orientations. To calculate this loss, output map 1 mentioned above is rotated computationally to produce output map 3. Then output map 4 is obtained by applying the 3DFSC filter to output map 3. This output map 4 is processed through the neural network to generate output map 5. Minimization of equivariance loss ensures that the output map 5 matches output map 3.

The noise2noise consistency and noise2noise equivariance losses are similar to consistency and equivariance losses, respectively. Recent cryoEM and cryo electron tomography (cryoET) packages, such as Cryo-CARE^36^, Topaz-denoise^37^, WARP/M^8, 38^, and RELION5^39^, have implemented denoising convolutional neural network in the absence of explicit access to ground-truth images with the noise2noise framework^35^. Noise2noise relies on pairs of noisy images to extract information about their shared signal. In the spIsoNet implementation, the output maps generated from two independent determined 3D half-maps are treated as a pair for noise2noise network training (Fig. 1c). Specifically, the noise2noise consistency loss calculates the difference between output map 2 and the submap extracted from the other half-map. To calculate the noise2noise equivariance loss, the submap from the other half-map is passed through the neural network to produce output map 1′, which is then rotated to produce output map 2′. The noise2noise equivariance loss is calculated to minimize the difference between output map 2′ and output map 5. This denoising algorithm acts as a local filter, improving the map quality and suppressing artifacts (Fig. 1d).

We tested the limit of information recovery for *Anisotropy Correction* with simulated data. A ribosome density map was created from the atomic model of a translating 70S ribosome (PDB-8B0X)^40^ and used to simulate distorted maps suffering from various levels of preferred orientation problem. We applied a missing-cone filter that zeros values in the Fourier space up to certain angles to create three different maps with missing cones, mimicking different severities of map anisotropy (Fig. 2a-i). Applying larger (45°, 60° and 75°) angles of the missing-cone filter results in more severe density elongation (Fig. 2b, e, h and Extended Data Fig. 2a-2c) and more distorted rRNA (Fig. 2c, f, i). Applying *anisotropy correction* improved the isotropy of all the three distorted maps, especially for the 45° and 60° missing-cone filtered maps, where elongation is eliminated and both bases and phosphates of nucleotides are well-resolved (Fig. 2b, c, e, and f). Even for the severe situation with −75° to 75° missing cones, spIsoNet was able to reduce elongation distortion to the RNA backbone (Fig. 2i and Extended Data Fig. 2a). We conclude that spIsoNet’s *anisotropy correction* module can faithfully and effectively recover missing information for simulated data suffering from a −60° to 60° missing cones.

**Figure. 2.**
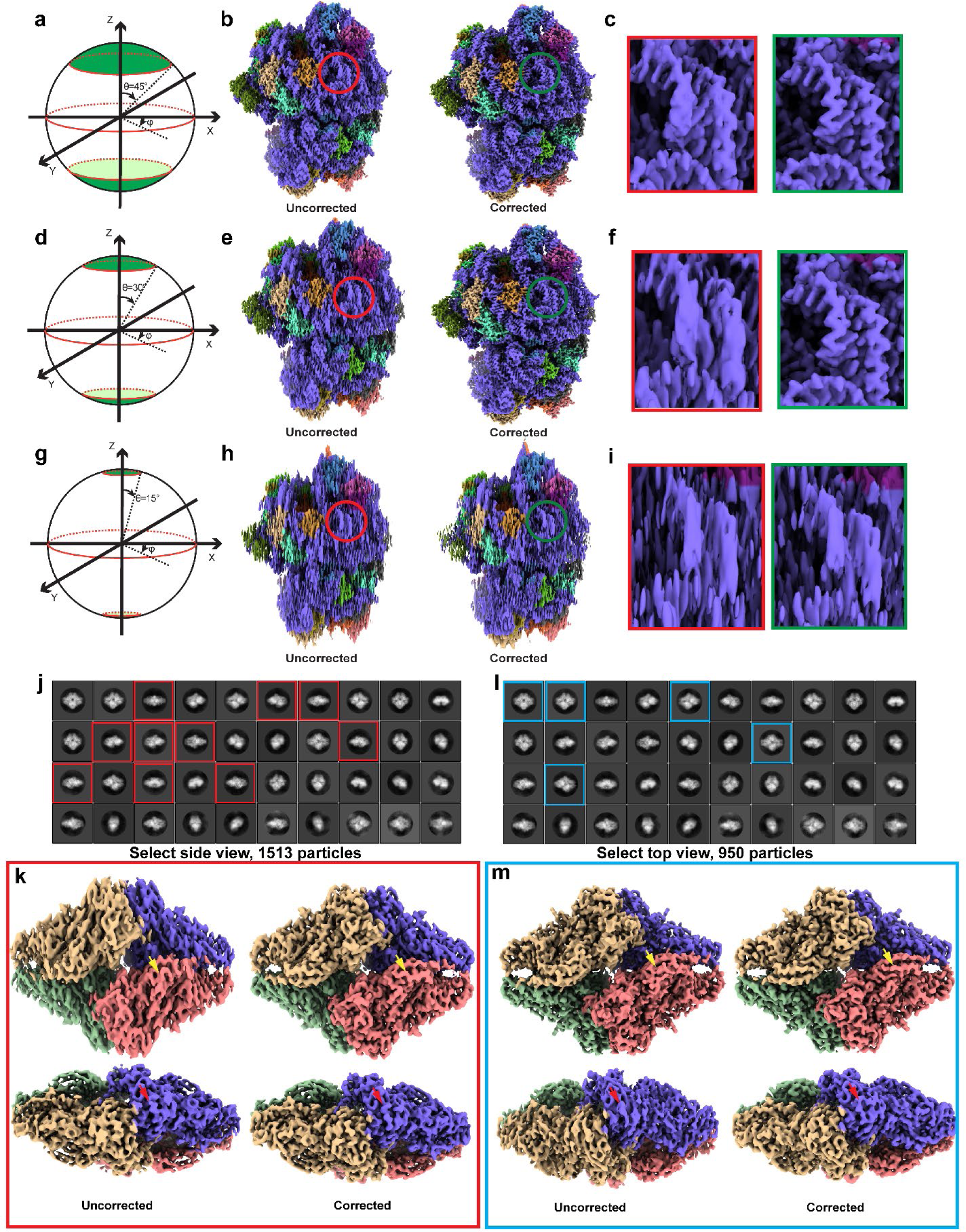
| Evaluation of information recovery by spIsoNet. **a-i,** *Anisotropy correction* applied to synthetic data generated from the ribosome structure (PDB-8b0x)^40^. Three different missing cone datasets were made for comparison. The red panel show zoom-in the density regions of the uncorrected map, while the green panel show the density regions of corrected map after *Anisotropy Correction*. **j-m,** *Anisotropy Correction* applied to cryoEM β-galactosidase density maps reconstructed from different views in 2D classifications in **j** and **l**.

### The Misalignment Correction module

The *Misalignment Correction* module is an integrated workflow comprising map filtering, anisotropy correction, and RELION auto-refine (Fig. 1e). The results of this module include improved particle orientation parameters (saved in STAR file) and two reconstructed half-maps from RELION. After each iteration of 3D refinement, post-processing-like filters including whitening^41^ and FSC weighting^42^ are applied to the produced half-maps. Then, spIsoNet’s *Anisotropy Correction* module is applied to these filtered half-maps. The resulting corrected half-maps are then low-pass filtered to their corresponding resolution based on the gold-standard FSC 0.143 criterion and the two filtered half-maps are used as reference for the next iteration of orientation estimation independently. This module has an option “--keep_lowres” to preserve the reliable low resolution (10-20 Å) information from the initial reference map to enhance accuracy of the initial alignment. This option is desirable for specimens that suffer from severe preferred orientation where a correct initial map could not be obtained. Though low-resolution information from the reference is used for alignment, the final reconstruction is obtained solely from original particles.

Notably in this module, the neural network used in spIsoNet exploits prior knowledge (consistency and equivariance losses) to regularize cryoEM reconstruction. This practice of using deep-learning to regularize the structure determination process is akin to the denoising algorithms implemented in M^8^ or RELION5 Blush regularization^39^. But unlike these methods, we not only have the denoiser neural network, but also incorporate the isotropic prior information to further improve the resolution and density quality of the reconstructed maps.

### Applications of spIsoNet in real-world preferred orientation problems

#### Application of the Anisotropy Correction module to a tetrameric protein complex—the β-galactosidase

We tested the spIsoNet on an RELION tutorial dataset containing β-galactosidase. By selecting side-view particles and top-view particles from 2D class averages (Figure 2j, l), we curated two subsets of particles with preferred orientation and performed standard RELION 3D reconstructions (Fig. 2k, m). Because accurate alignment of the particles is already available, we only apply *Anisotropy Correction* to obtain corrected maps (Fig. 2k, m). The corrected maps have reduced density elongation, continuous β-strands, and clearer α-helical pitch. Hence, our tests on the β-galactosidase dataset demonstrate that the *Anisotropy Correction* module alone robustly reduces artifacts caused by top-view dominant or side-view dominant orientations.

#### Application to a protein dataset with moderate preferred orientation problem— hemagglutinin trimer with 40° tilt

The cryoEM map (EMDB-8731) of HA trimer from dataset (EMPIAR-10097) acquired at 40° stage tilt in previous study^10^ shows relatively poor quality as illustrated by unclear side chain densities like rod-like α-helices (Fig. 3a-b). To evaluate the performance of spIsoNet, we first performed *Anisotropic Correction* on the maps and discovered that the corrected map exhibits better quality with higher local resolution and less noise (Extended Fig. 3a and 4b). The side chain densities which cannot be observed in the original map are discernable in the anisotropy-corrected map (Fig. 3a-b). We then applied *Misalignment Correction* module on this particle dataset and obtained a cryoEM map at 4.1 Å resolution (Fig. 3a) with well-resolved α-helical pitch, clear side chain densities (Fig. 3b), and near-spherical (0.991) 3DFSC (Fig. 3c, e). The misalignment-corrected map shows improved half-map FSC and map-to-model FSC (Extended Data Fig. 3b-c), as well as larger isotropic regions of Fourier shell occupancy^43^ (FSO; Fig. 3d, f) compared with the original map. Therefore, for this dataset, both *Anisotropic Correction* and *Misalignment Correction* significantly improves the quality of cryoEM reconstructions.

**Figure. 3.**
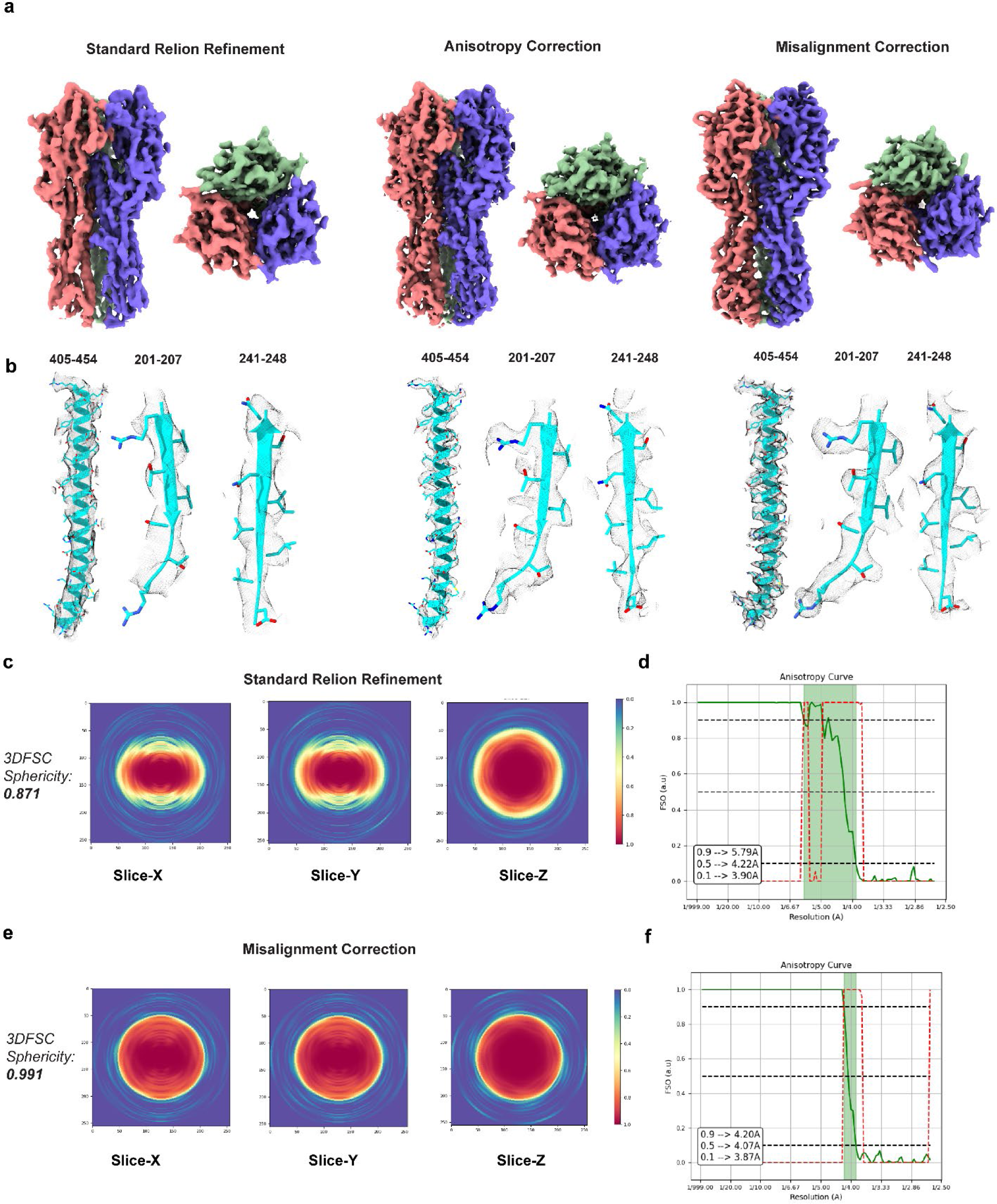
| The application of spIsoNet to the tilted cryoEM influenza hemagglutinin (HA) trimer datasets. **a,** cryoEM map of the HA trimer reconstructed from different methods. From left to right: standard RELION Refinement, *Anisotropy Correction*, and *Misalignment Correction*. **b,** Representative density regions of the HA trimer structure with the HA atomic model (cyan) fitted. The residue numbers are indicated. From left to right: standard Relion Refinement, *Anisotropy Correction*, and *Misalignment Correction*. **c,** Central slices along the X, Y and Z directions of the 3DFSC for the standard RELION Refinement. **d,** FSO (green line) and P value for the Bingham test (red dashed line) computed from the standard RELION Refinement result. **e,** Central slices along the X, Y and Z directions of the 3DFSC for the spIsoNet misalignment corrected result. f, FSO (green line) and P value for the Bingham test (red dashed line) computed from the spIsoNet misalignment corrected result.

**Figure. 4.**
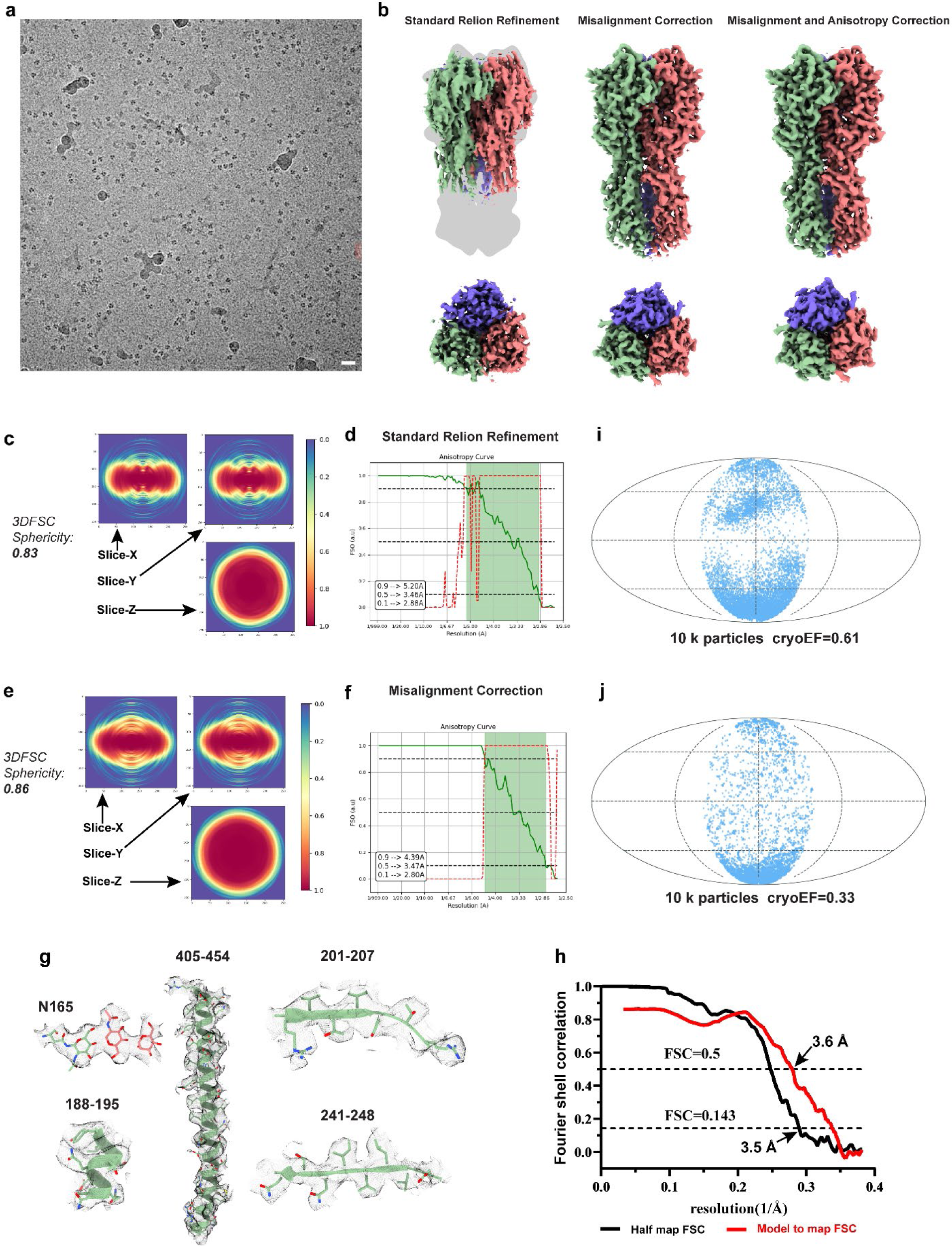
| The application of spIsoNet to the untilted cryoEM influenza hemagglutinin (HA) trimer datasets. **a,** Representative cryo-EM micrograph of the untilted cryo-EM influenza HA trimer dataset (scale bar, 20 nm). **b,** cryoEM map of the HA trimer reconstructed from different methods. From left to right: standard RELION Refinement, spIsoNet *Misalignment Correction*, and both *Misalignment and Anisotropy Correction*. **c,** Central slices along the X, Y and Z directions of the 3DFSC for the standard RELION Refinement. **d,** FSO (green line) and P value for the Bingham test (red dashed line) computed from the standard RELION Refinement result. **e,** Central slices along the X, Y and Z directions of the 3DFSC for the spIsoNet misalignment corrected result. **f,** FSO (green line) and P value for the Bingham test (red dashed line) computed from the spIsoNet misalignment correction result. **g,** Representative density of selected amino acid residues and glycans (denoted with the residue numbers) from the misalignment corrected HA timer cryoEM map docked with the relevant atomic models. **h,** The Fourier shell correlation (FSC) curves of the misalignment corrected HA trimer map. Black, half-map FSC curve with criterion of 0.143; Red, FSC curve calculated between the cryo-EM map and the refined structure model with criterion of 0.5. **i,** The orientation distribution of standard RELION Refinement result created by the cryoEF. **j,** The orientation distribution of spIsoNet misalignment corrected result generated by the cryoEF.

#### Application to a protein dataset with severe preferred orientation problem— hemagglutinin trimer without tilt

The untilted dataset (EMPIAR-10096) of HA trimer experiences severe preferred orientation (Fig. 4a) leading to failure of generating a trustworthy cryoEM map^10^ using standard cryoEM structure determination pipeline. This cryoEM map is much shorter than the true structure and without interpretable secondary structure (Fig. 4b). Since spIsoNet’s *Anisotropy Correction* relies on reasonably accurate information within the cryoEM map, this erroneous structure does not benefit from the *Anisotropy Correction*. Instead, we performed the spIsoNet’s *Misalignment Correction* on the particle datasets, using the HA trimer map reconstructed from the tilt dataset as reference (Fig. 3a, right panel). During every iteration of refinement, we kept low resolution information up-to 10 Å from the reference for reliable orientation alignment. With *Misalignment Correction*, we obtained a reconstructed map with correct shape (Fig. 4b) and improved map isotropy (Fig. 4c-f). We observed discernable amino-acid side chains and glycans in the map sufficient for atomic model building (Fig. 4g), consistent with the resolution determined by half-map (3.5 Å) and model-to-map (3.6 Å) FSC (Fig. 4h). Noteworthy, for standard RELION reconstruction, the cryoEF score^44^ are exaggerated because of the existence of misaligned particles (Fig. 4i). After *Misalignment Correction*, the particle orientation distribution shifted and results in a more reasonable cryoEF score reflecting a true severe preferred orientation (Fig. 4j). This particle misalignment-corrected map can be then treated by *Anisotropy Correction*, further leading to a denoised structure with more continuous densities (Fig.4b, Extended Data Fig. 5).

#### Application to a dataset of an asymmetric, protein-nucleic acid complex

Using the *Acinetobacter baumannii* 70S ribosome dataset (EMPIAR-10406)^45^, we curated a preferential-oriented dataset, by choosing particles within a specified Euler angles range [rot angles (−20°– 140°) and tilt angles (100°–160°)]. Standard reconstruction using these particles yields a map exhibiting elongated and fragmented densities (Fig. 5a-b and Extended Data Fig. 6a). With *Anisotropic Correction*, the corrected map showed significant improvement in the map quality, represented by continuous map density, higher local resolution, and less noise (Fig. 5a-d and Extended Data Fig. 6b, 7a-b). We then performed both *Misalignment and Anisotropic Correction* on this dataset. We observed that this combined protocol improved the reconstruction with further enhanced visibility of side chains (Fig. 5a-d). We observed that utilizing subtomogram averaging maps from either 70S or 80S ribosomes (Extended Data Fig. 7c-d) as references, while maintaining a 15Å initial resolution for alignment (Extended Data Fig. 7e-f), consistently yielded comparable high-quality maps without model bias (Fig. 5a-d). Thus, spIsoNet improve particle alignment and mitigate anisotropy for asymmetric and nucleic acid-containing molecules.

**Figure. 5.**
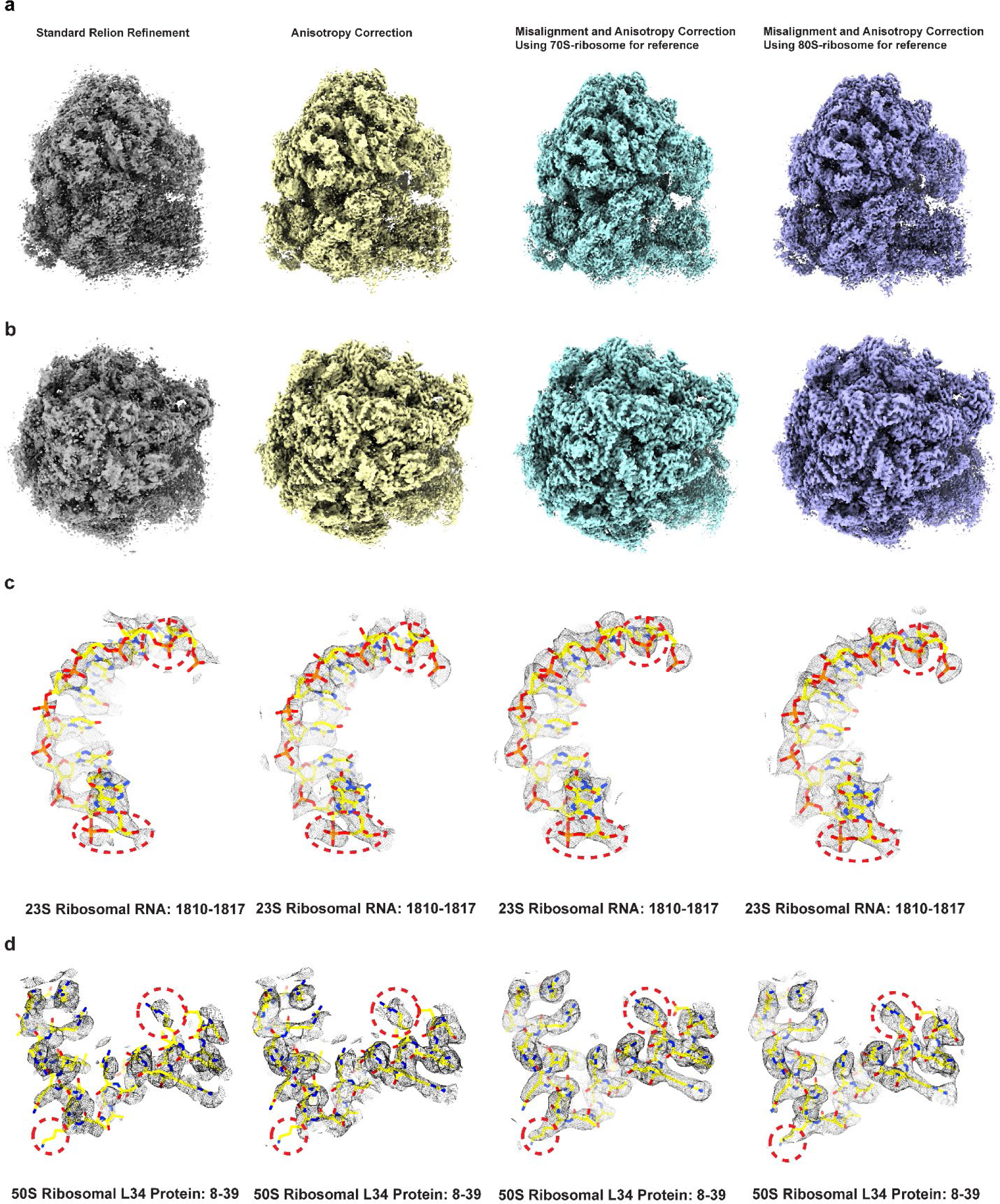
| Application of spIsoNet to the ribosome dataset. **a-b,** Ribosome maps reconstructed from different reconstruction methods. From left to right: standard RELION Refinement, spIsoNet *Anisotropy Correction*, *Misalignment* followed by *Anisotropy correction* using 70S ribosome as reference, and *Misalignment* followed by *Anisotropy Correction* using 80S ribosome map as reference. **c-d,** Representative density regions with the fitted atomic model (yellow).

### spIsoNet can also improve map quality and isotropic resolution in subtomogram averaging

CryoET is a method of choice for obtaining *in situ* biological structures by combining views in a tilt series of the same specimen area^46^. Bio-macromolecules are often arranged in specific ways in cells other than randomly oriented^47^, giving rise to preferred orientation problems *in situ* and distorted structures even after subtomogram averaging. We explored the performance of spIsoNet in subtomogram averaging, with the HIV-1 immature capsid dataset (EMPIAR-10164)^48^. Using a subset of five tilt-series, standard workflow in RELION4^49^ yields a 3.7 Å resolution structure without performing CTF refinement and frame alignment (Fig. 6a, Extended Data Fig. 8a). With *Misalignment Correction*, we obtained an isotropic 3.6 Å resolution structure (Fig. 6e, Extended Data Fig. 8a). Despite only marginal improvement in the overall resolution, the corrected map displayed improvements across various metrics including half-map FSC, model-to-map FSC, and local resolution (Fig. 6a, e and Extended Data Fig. 8). The structure revealed better resolved side chain densities (Fig, 6d and h), exhibited increased 3D FSC sphericity and reduced resolution anisotropy regions within the FSO curve (Fig. 6b, c, f, g). These improvements can be attributed not only to the information retrieval but also to the denoising effect, which should improve the accuracy of particle orientation estimation. Thus, spIsoNet can be used for *in situ* structural biology to improve map quality in subtomogram averaging.

**Figure. 6.**
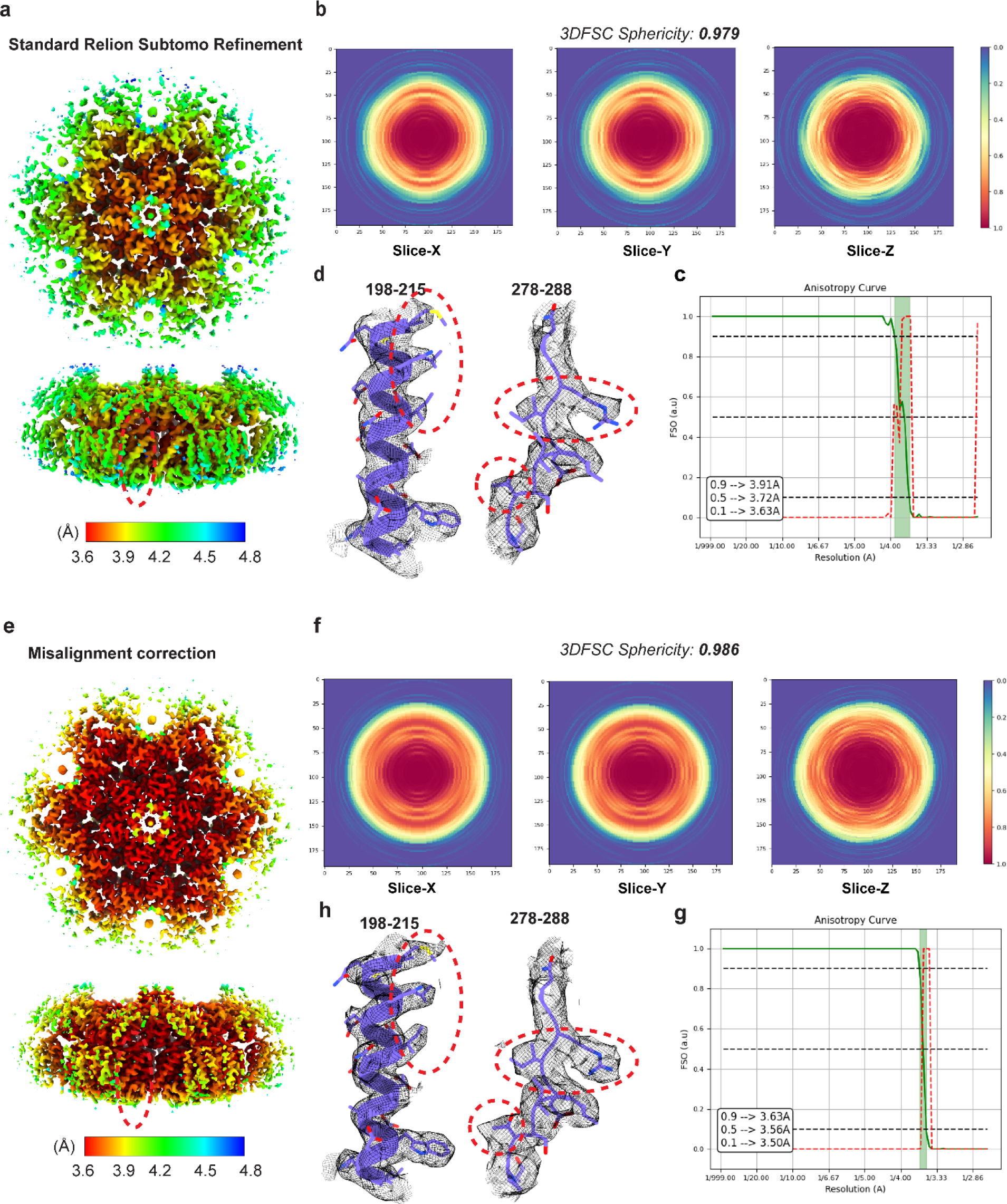
| The application of spIsoNet to the subtomogram-averaging. **a,** Local resolution map of the HIV-1 immature capsid reconstructed from standard RELION4 subtomogram-averaging. **b,** Central slices along the X, Y and Z directions of the 3DFSC for the standard RELION subtomogram averaging. **c,** FSO (green line) and P value for the BT (red dashed line) computed from the Standard RELION4 subtomogram averaging pipeline. **d,** Representative density of amino acid residues (denoted with the residue numbers) docked with the atomic models. **e,** Local resolution map of the HIV-1 immature capsid reconstructed from spIsoNet *Misalignment Correction*. **f,** Central slices along the X, Y and Z directions of the 3DFSC for the spIsoNet *Misalignment Correction*. **g,** FSO (green line) and P value for the Bingham test (red dashed line) computed from the spIsoNet result. **h,** Representative density of selected amino acid residues (denoted with the residue numbers) fitted with the atomic models.

## Discussion

In contrast to the numerous previous chemical and physical methods to alleviate the seemly ubiquitous problem of preferred orientation in cryoEM, spIsoNet now provides a computational method to address the preferred orientation problem. While the two modules in spIsoNet can be executed independently, using the *Misalignment Correction* module followed by the *Anisotropy Correction* module tends to yield optimal results (Fig. 3-5). For the severely preferred orientated particles, the *Misalignment Correction* module has an option of preserving the low-resolution information from the initial reference map to ensure the alignment accuracy (Fig. 4). The reference map can be obtained from various methods such as reconstruction from small particle datasets with stage tilt (Fig. 3) or subtomogram averaging from small tomography datasets (Fig. 5). Regardless which of the suggested spIsoNet workflows (Fig. 1f) users choose to solve their specific preferred orientation problem, the pure computational nature of the workflow should significantly shorten the imaging to structure interpretation, allowing investigators to focus on scientific discovery other than specimen/grid manipulation as needed for previous methods.

It is important to note that spIsoNet differs completely from IsoNet^50^ in both applications and algorithms. Self-supervised information recovery was also implemented in the original IsoNet package, which was designed to overcome the missing-wedge problem in electron tomography^50^. There, the network is trained with rotated subtomograms as ground truth, and rotated-and-missing-wedge-imposed subtomograms as input. As detailed above, due to the complexity of preferred orientation problem in cryoEM, here we had to use four-loss end-to-end implementation and 3DFSC as reliability measure in spIsoNet, thus different from the iterative one-loss design of IsoNet^50^.

Previous deep learning methods, such as deepEMhancer^51^ and EMReady^52^, have attempted to enhance the interpretability of cryoEM maps by training a network using maps simulated from the PDB as ground truth and cryoEM maps in the EMDB as input. While these approaches have shown promising results, concerns exist regarding to potential biases introduced by relying on knowledge from other structures, which may complicate the interpretation of neural-network-generated cryoEM maps. By contrast, spIsoNet exploits the rotation equivariance present in the biological structure with self-supervised learning. In essence, a cryoEM map contains molecular information distributed across a 3D volume, encompassing recurring information, ranging from large structural features like α-helices and β-sheets to small structural features in individual amino acids. By accounting for both consistency and equivariance loss in the network training process, the network learns how to recover information, leveraging the fact that characteristic features should be similar regardless of their positions and orientations, and ensures that the recovered molecular information will go to regions where information is lacking. The spIsoNet’s ability to capture complex patterns and relationships solely from user-provided cryoEM maps allows it to achieve faithful representations of reality with incomplete experimental data.

In single particle cryoEM, the accuracy of reconstruction hinges upon how to incorporate reliable prior knowledge into the cryoEM structure determination pipeline. From smoothness priors in RELION^5^, to the independent half-map noise assumptions^39, 53^, to low-dimensional manifold of conformation change^54^, and to the isotropic prior in spIsoNet, reliable prior knowledge is incorporated into the reconstruction process. Such integration allows for 3D reconstructions from fewer particles and less orientation views, enhancing the accuracy, efficiency, and investigative power of cryoEM and cryoET in biological inquiries.

## Methods

### Software implementation

We implemented spIsoNet in Python using Linux as the native operating system. The package can be downloaded from Github (https://github.com/IsoNet-cryoET/spIsoNet). A detailed document is provided, accompanied by the spIsoNet package.

### Overview of Anisotropy Correction module

The input of spIsoNet’s *Anisotropy Correction* are two unfiltered half-maps and a soft solvent mask. It first calculates 3DFSC volume using the two half-maps to represent the anisotropic resolution for the reconstruction. Then IsoNet perform the refine step to train a network to recover the missing information quantified based on 3DFSC^10^. The refine step will perform on the half-maps, generating a neural network and the anisotropy corrected half-maps (Fig. 1c, Extended Data Fig. 1). The subsequent postprocessing of the half-maps for sharpening and gold-standard FSC determination can be performed in other cryoEM softwares such as RELION^5^.

### Fast 3DFSC Implementation

3DFSC^10^ is an algorithm that quantitatively assesses directional resolution anisotropy of density maps. The algorithm compiles a set of 1D Fourier shell correlation curves computed over distinct angular directions into an 3D array, which is visualized as a 3D density volume.

For user convenience and to make our program standalone, we implemented the 3DFSC algorithm in the spIsoNet natively. We used a cKDTree algorithm^56^ to increase the speed of processing. It typically runs for 30 second for maps with 400^3^ voxels on 16 CPU cores.

In the following sessions, applying the 3DFSC filter to a 3D volume x is denoted as Ax, which means applying this filter in Fourier space, to mimic the preferred orientation induced artifacts on map x:

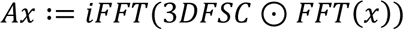

Where FFT and iFFT are 3D Fourier transform and 3D inverse Fourier transform respectively. ⊙ is dot product.

### The neural network architecture in spIsoNet

The neural network used in spIsoNet is based on U-net^33^ (Fig. 1b), which is well recognized in biomedical image restoration and segmentation. The main building blocks of our U-net implementation is a 3D convolution layers with non-linear activation functions called leaky Rectified Linear Units, which are applied per voxel. Those convolution layers have kernel sizes of 3 × 3 × 3. Three 3D convolution layers are stacked together to form a convolution block in our network, which can extract features in cryoEM maps. By stacking the convolutional blocks, the U-net is built based on encoder-decoder architecture (Fig. 1b). The encoder path is a set of convolution blocks and strided convolution layers that compress 3D volumes. Strided convolution layers reduce the spatial size of the input of this layer by 2 × 2 × 2, allowing the network to learn more abstract information. one encoder block is composed of a convolution block followed by a strided convolution layer. Three encoder blocks constitute the entire encoding path. The number of convolution kernels for each convolution layer doubles after each encoder block. After the encoder path, the 3D volumes are processed with a convolution block and enter the decoder path of the network. The decoder path is symmetrical to the encoder but uses transpose convolution layers, opposite to strided convolution layers, to enlarge the dimension of features. Although the down-sampling of the 3D volumes captures the essence of the features, high-resolution information is lost by stride convolution operations. In U-net, skip-connections that concatenate the feature layers of the same dimension in two paths are implemented to preserve high-resolution information.

In the following sessions the network is represented as *f*_*θ*_(), which captures the neural network as f with parameters θ. The parameter parameters θ are network weights and biases that are updated throughout training process.

### Network training for *Anisotropy Correction*

Firstly, voxels values of input half-maps are normalized to standard deviation of 1 and mean of 0. By default, 1000 randomly distributed 3D coordinates are randomly generated within the regions defined by a user-provided mask. Then, cubic volumes (default 64^3^ voxels) centered at the generated coordinates are saved as submaps. The network is trained to minimize a loss function. The loss function is minimized by employing Adam optimizer with an initial learning rate of 0.0004. The neural network training in is performed on one or multiple GPUs and typically consists of 30 epochs.

The loss function consists of four terms: two for information recovery and two for noise2noise based denoising.

The first term is the data consistency loss:

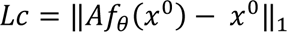

In this formula, *x*^0^ represents the input submaps in one half-map, *A* denotes the 3DFSC filter described above, and *f*_*θ*_ represents the neural network parameterized by θ. This term in this expression enforces data consistency, ensuring that the application of the 3DFSC filter to the predicted map results in the raw map.

The second term is the rotation equivariance loss:

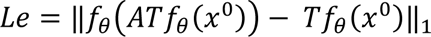

This loss takes the rotated map and tends to recover information from the removed information in the rotated maps. T stands for random rotation operator. To calculate the rotation equivariance loss, we create a rotated and corrected submap (R map, *i.e. Tf*_*θ*_(*x*)) by applying the network to an input submap and randomly rotating the resulting map. After applying the 3DFSC filter to the R map, we pass it through the network again for recovery. The rotation equivariance terms minimize the difference between the recovered map and the original R map.

The third and the fourth terms are similar data consistency loss and rotation equivariance loss but comparing between the submaps from half-map1 and half-map2 for the noise2nosise based denoising:

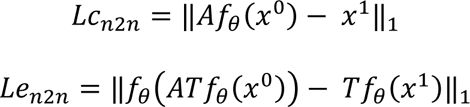

Where in these terms *x*^0^ and *x*^1^ are submaps from different half-maps, but at the same location. Noise will be reduced using these two terms which benefits spIsoNet information recovery.

The final loss is represented as:

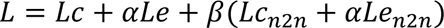

Where *α* and *β* are hyperparameters. The *α* is balancing between data consistency and rotation equivariance, and beta defines how much level of denoising should be considered in training. By default, *α* and *β* values equal to 1 and 0.5, respectively.

After the last epoch of training. the spIsoNet program splits the half-maps into smaller 3D chunks and applies the network to them separately. Then output 3D chunks are stitched to produce the final output. To avoid the line artifact between adjacent chunks caused by the loss of information on the edges of subtomograms. We implemented a seamless reconstruction method called overlap-tile strategy^33^, which predicts the overlapping chunks to avoid the edge effect. The output corrected two half-maps as well as the trained neural network are saved for further post-processing.

### Particle Misalignment Correction module

In the particle *Misalignment Correction* module, we integrate spIsoNet in RELION’s 3D refinement pipeline. Taking the advantage of “--external_reconstruct” function in *relion_refine*, the *relion_wrapper.py* in spIsoNet will be executed after each iteration of RELION refinement when RELION refinement reaches a finer angular sampling, *i.e.* “healpix order” becomes 3.

After each RELION iteration, the spIsoNet will be executed in 4 steps: spectral whitening, FSC weighting, *Anisotropy Correction*, and low-pass filtering. Spectral whitening equalized the Fourier power of half-maps for all the spectral frequency from 10Å up to the resolution determined by FSC=0.143 criterion, similar to map whitening implemented in cisTEM. Those maps are then performed FSC weighting. Because the FSC weighting is performed only on the half-maps, the 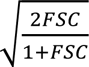 weighting proposed in Rosenthal and Henderson is not performed^42^, instead the half-maps are directly filtered with the FSC. After the FSC weighting, map *anisotropy correction* is performed for those half-maps. Those maps are then low-pass filtered to a resolution of FSC=0.143 and serve as references for the subsequent iterations. Users can choose not to perform either spectral whitening, FSC weighting, or low pass filtering, if necessary.

In some cases, due to the severe preferred orientation problems, RELION fails to perform reliable image alignment and to generate relatively correct initial maps. As a result, we provided an option called “--keep_lowres”. This option allows for preserving the low-resolution information of the reference maps during the subsequent RELION refinement iterations. Given a relatively uniformly sampled reference at about 10-20 Å resolution, RELION will be less likely to perform erroneous image alignment. The low-resolution reference map can be obtained from various methods such as reconstruction from small particle datasets with stage tilt (Fig. 3), or subtomogram averaging from a small tomography dataset (Fig. 5). To avoid differences in the intensity between the reference map and the map reconstructed from the particle dataset that could cause errors in alignment, we designed a function called spectral matching that enforces the spectral power distribution of the reference maps the same as the map reconstructed using the particle datasets in each RELION iteration. This “keep_lowres” step is performed before the whitening step.

### Benchmark with simulated ribosome map

To benchmark spIsoNet’s performance with an asymmetric complex, we downloaded the structure of the a translating 70S ribosome (PDB-8B0X)^40^ and generated a density map with a pixel size of 1 Å and a box size of 400×400×400 from the atomic model. We then applied a missing cone-filter that remove orientational information in Fourier space up to certain degrees to create three different maps with missing cones (Fig. 2a, d, g), mimicking different degrees of map anisotropy conditions in practice. Because we only have one map for each simulation, we cannot calculate 3DFSC. Instead, we used missing cone volumes as a quantification of the missing information in Fourier space. The spIsoNet α parameter is set to 1. Also, because there is only one map, β parameter is not applicable in this experiment. Total 30 epochs are performed for the correction of each simulated map.

### Benchmark with Beta-galactosidase datasets

We also tested spIsoNet with the β-galactosidase dataset which was used as the RELION single particle tutorial dataset. To validate whether spIsoNet can improve the map quality caused by different views of preferred orientation, we created two different datasets with preferred orientation by selecting side view particles (1,513 particles) and top view particles (950 particles) from the 2D class averages (Fig. 2j-m).

We conducted standard RELION 3D reconstruction to generate two anisotropic maps, as depicted in (Fig. 2k and 2m). Subsequently, we applied map *Anisotropy Correction* to obtain the corrected maps, as shown in (Fig. 2k and 2m). The *Anisotropy Correction* was executed with parameters set to α=1, β=0.5, limiting information recovery resolution up-to 3.5 Å, and running for 30 epochs. Both side-view dominant and to-view dominant maps corrected by anisotropy correction shows improved map quality with significantly reduced the density smearing and elongation on the missing information direction.

### Benchmark with the HA-trimer tilted dataset: EMDB 8731 and EMPIAR-10097

We first downloaded the 4.2 Å HA trimer cryoEM map (EMDB-8731)^10^ and performed *Anisotropy Correction* directly. The *Anisotropy Correction* parameters are α=1 and β=0.5, with limit information recovery resolution to 3.5Å. After *Anisotropy Correction*, Postprocessing is performed using RELION with default parameters for structure comparison.

The EMPIAR-10097 dataset of HA-trimer was collected using grid-tilting strategy^10^. We imported 13,0000 particles into RELION4 and performed *Misalignment Correction*, when the healpix-order became 3. We set the SpIsoNet parameters as follows: α=1, β=0.5, and epochs=5. We retrained the spIsoNet model with every RELION4 refinement iteration using the RELION generated half-maps. We finally obtained the 4.1 Å resolution map measured by gold standard FSC.

### Benchmark with HA-trimer non-tilted dataset: EMPIAR-10096

This cryoEM dataset of HA-trimer was collected without grid tilting^10^. 13,000 particles were imported into RELION4 and performed the *Misalignment Correction* using “--external_reconstruction” and “--keep_lowres” command when the healpix-order became 4. The “--keep_lowres” command allows us to incorporate the low-resolution initial map into each round of RELION4 reconstruction to enhance the initial 3D alignment accuracy. The initial reference is the map reconstructed from the tilt dataset (Fig. 3a, right panel) and filtered to 10 Å resolution. The spIsoNet *Misalignment Correction* parameters were set as follows: α=1, β=0.5, epochs=5. We retrained the spIsoNet model with every RELION4 refinement iteration. After RELION refinement, we suspect that there are some bad particles. Thus, we performed 3D classification without alignment for 4 classes. We selected two of the resulting classes with total 85,358 particles for another round of the *Misalignment Correction* with the same parameter as before. We finally obtained a 3.45 Å resolution map. We further performed the *Anisotropy Correction* to improve the cryoEM map. The parameters of *Anisotropy Correction* are α=1, β=0.5, epochs=30 and limit recovery resolution to 3.5Å.

### Benchmark with asymmetric ribosome dataset: EMPIAR-10406

This dataset contains the 70S ribosome from the human pathogen *Acinetobacter baumannii* in complex with amikacin^45^. We manually created a preferred orientation datasets by selecting the particles with specific ranges of Euler angles: rot angles (−20°–140°) and tilt angles (100°–160°). We followed the standard single particle RELION4 protocol to perform the 3D auto-refinement using 23,177 particles. We then executed the *Misalignment Correction* using spIsoNet with “--external_reconstruction” and “--keep_lowres” commands when the healpix-order became 3, the initial resolution for the reference is 15 Å and the information lower than this resolution is maintained through the intermediate RELION refinement process. The spIsoNet parameters were set as follows: α=1, β=0.5, epochs=5. Neural networks in spIsoNet are retrained after every RELION refinement iteration to adapt to the updated half-maps. To test the effect of incorporating the low-resolution information from the initial reference map, we chose two different subtomogram-averaging 3D cryo-EM maps for comparison: one is the 70S Ribosome structure from *E.coli* (EMDB-16139)^57^ and another is the 80S Ribosome structure from the *S.cerevisiae*.

To generate a reference map from the 80S Ribosome, we reprocessed the EMPIAR-10045 datasets^58^ as follows. The *S.cerevisiae* 80S Ribosome tilt-series were aligned using IMOD^59^, and the defocus values for each tilt were estimated using GCTF^60^. We picked 3,288 ribosome particles using TomoTwin software^61^ and performed the subtomogram averaging using RELION4^49^. We followed the standard RELION4 subtomogram refinement procedure and achieved a 7.23 Å resolution of yeast 80S ribosome.

### Benchmark with HIV VLP tomography dataset: EMPIAR-10164

This dataset contains the immature HIV-1 dMACANC VLPs^48^, which is also the RELION4 subtomogram averaging tutorial dataset^49^. We followed the standard RELION4 refinement procedure according to the tutorial and achieved a 3.7 Å resolution map. We then used the same RELION4 parameters and executed the *misalignment correction* with --external_reconstruction when the healpix-order became 4. The spIsoNet parameters were set as follows: α=1, β=0.5, epochs=20. We retrained the spIsoNet model with every RELION refinement iteration. We finally obtained a 3.6 Å resolution map. We used the unsharpened maps for comparison between the standard RELION refinement and *Misalignment Correction*.

### 3D visualization

IMOD^59^ was used to visualize 2D slices of cryo-EM density maps. UCSF ChimeraX^62^ was used to visualize the reconstructed cryo-EM density maps in their three dimensions. The 3DFSC and FSO are calculated within the Scipion software^63^. The local resolution maps are calculated by ResMap^64^ and local resolution estimation in RELION^5^.

## Data availability

The data that support this study are available from the corresponding authors upon request. The cryo-EM density maps of HA-trimer from EMPIAR-10096 and EMPIAR-10097 have been deposited in the EMDB under the accession numbers EMD-XXXXX and EMD-XXXXX, respectively.

## Code availability

The spIsoNet is available at the GitHub (https://github.com/IsoNet-cryoET/spIsoNet).

## Acknowledgements

This project is supported by a grant from the US National Institutes of Health (R01GM071940 to Z.H.Z.). We thank Pengcheng Zhou for helpful discussion.

## Author contributions

Y.-T.L. conceptualized the method, wrote code, processed data, made illustrations and documentation, and wrote the paper; H.F. comes up with the misalignment process, processed data, made documentation and illustrations, and wrote the paper; J.J.H. assisted in data processing, figure generation, and paper writing; Z.H.Z. oversaw the project, interpreted the results, and wrote the paper. All the authors edited and approved the manuscript.

## Reporting summary

Further information on research design is available in the Nature Research Reporting Summary.

## Extended Data Figures

**Extended Data Fig. 1.**
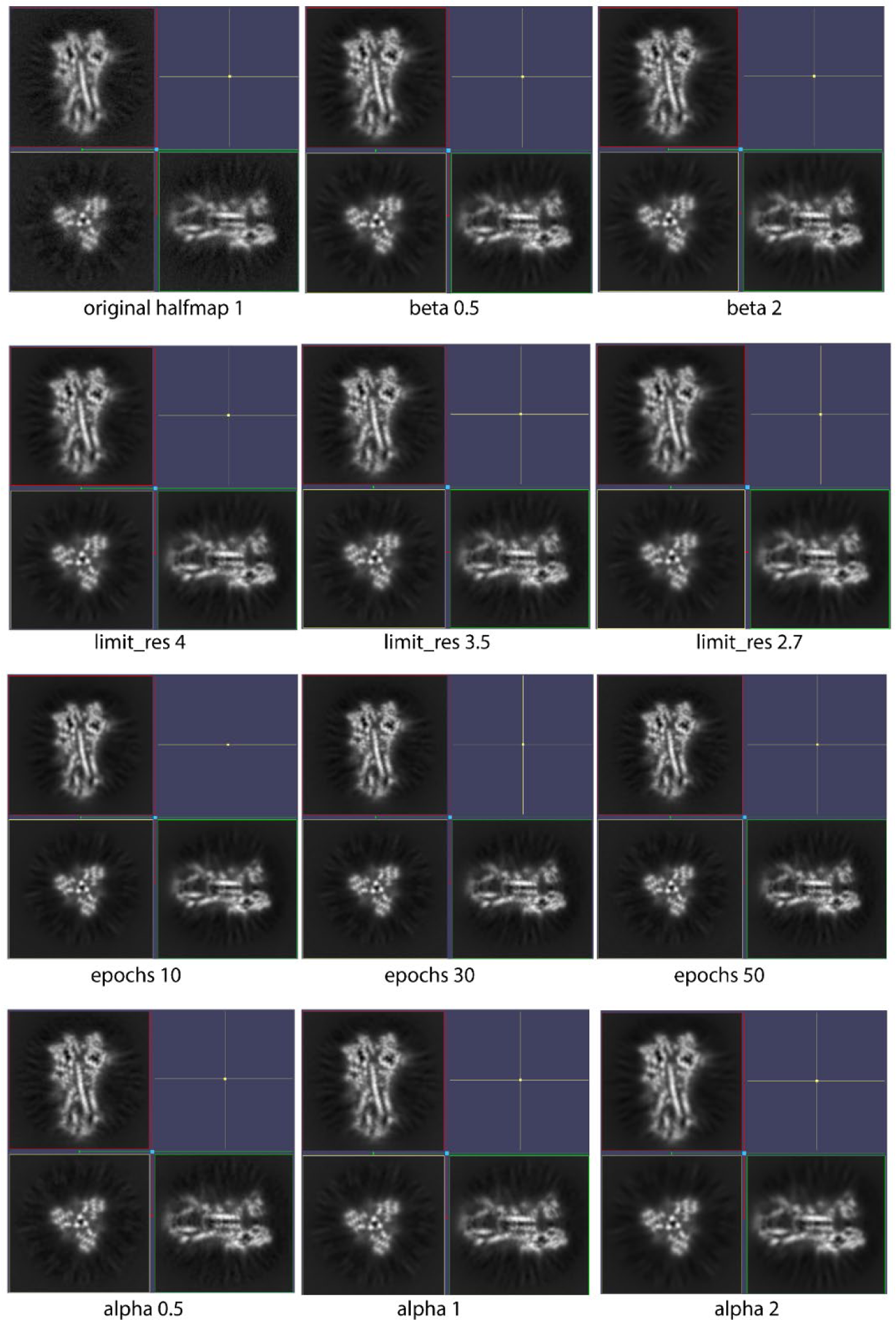
| The effect of the spIsoNet *Anisotropy Correction* on the HA with different α, β, epochs, and limit resolution values. Orthogonal slices of cryoEM half-map corrected using spIsoNet *Anisotropy Correction* with different parameters. The default parameters are α=1, β=0.5, epochs=30, limit_res=3.5. If the parameters are not indicated in the figure the default parameters are used.

**Extended Data Fig. 2.**
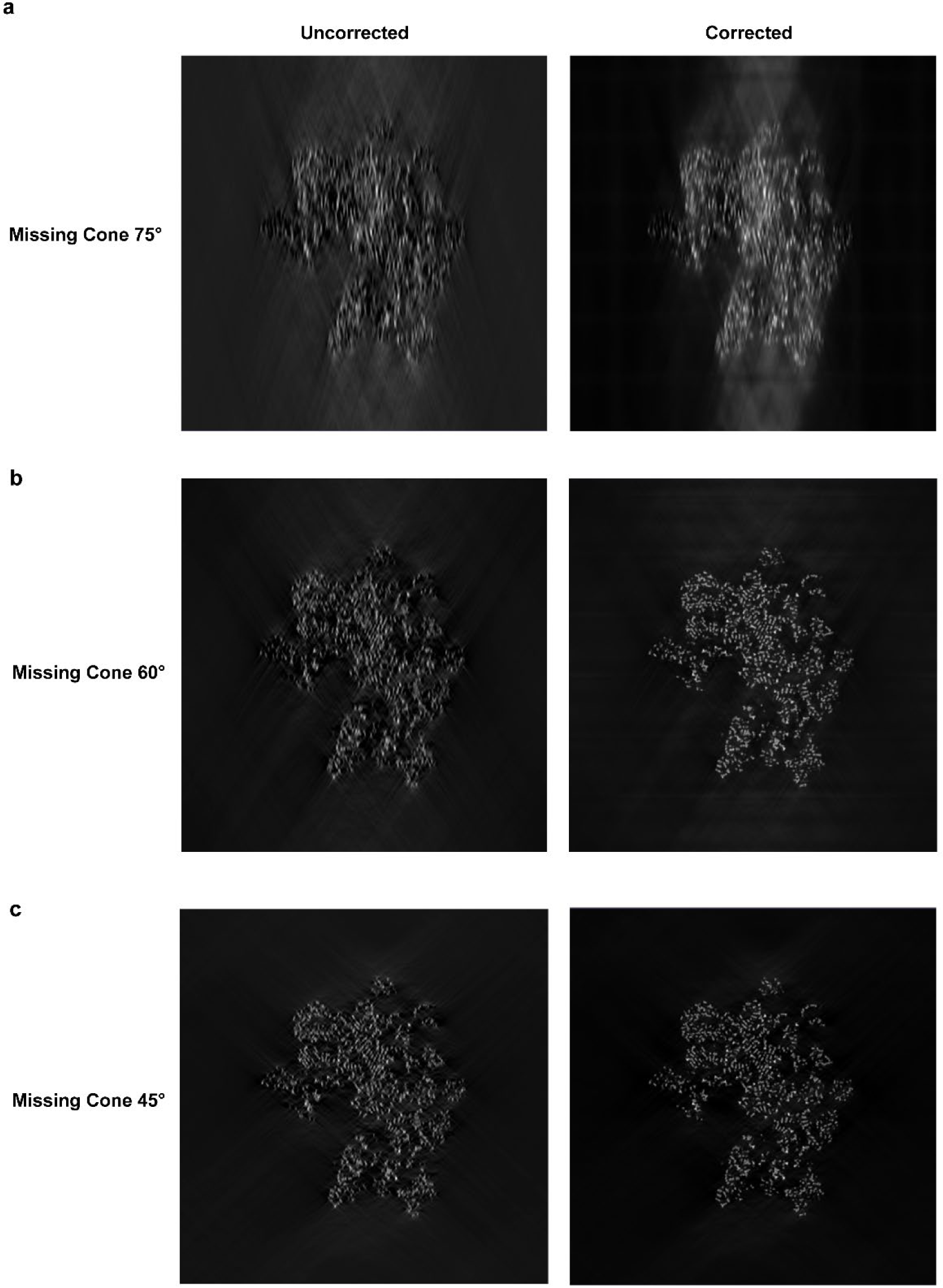
| The effect of the spIsoNet *Anisotropy Correction* on the simulated ribosome maps. **a-c,** Left panels show the sections of XZ planes for the uncorrected maps simulated with different missing cones, while the right panels show the sections on XZ planes for the maps with *Anisotropy Correction*.

**Extended Data Fig. 3.**
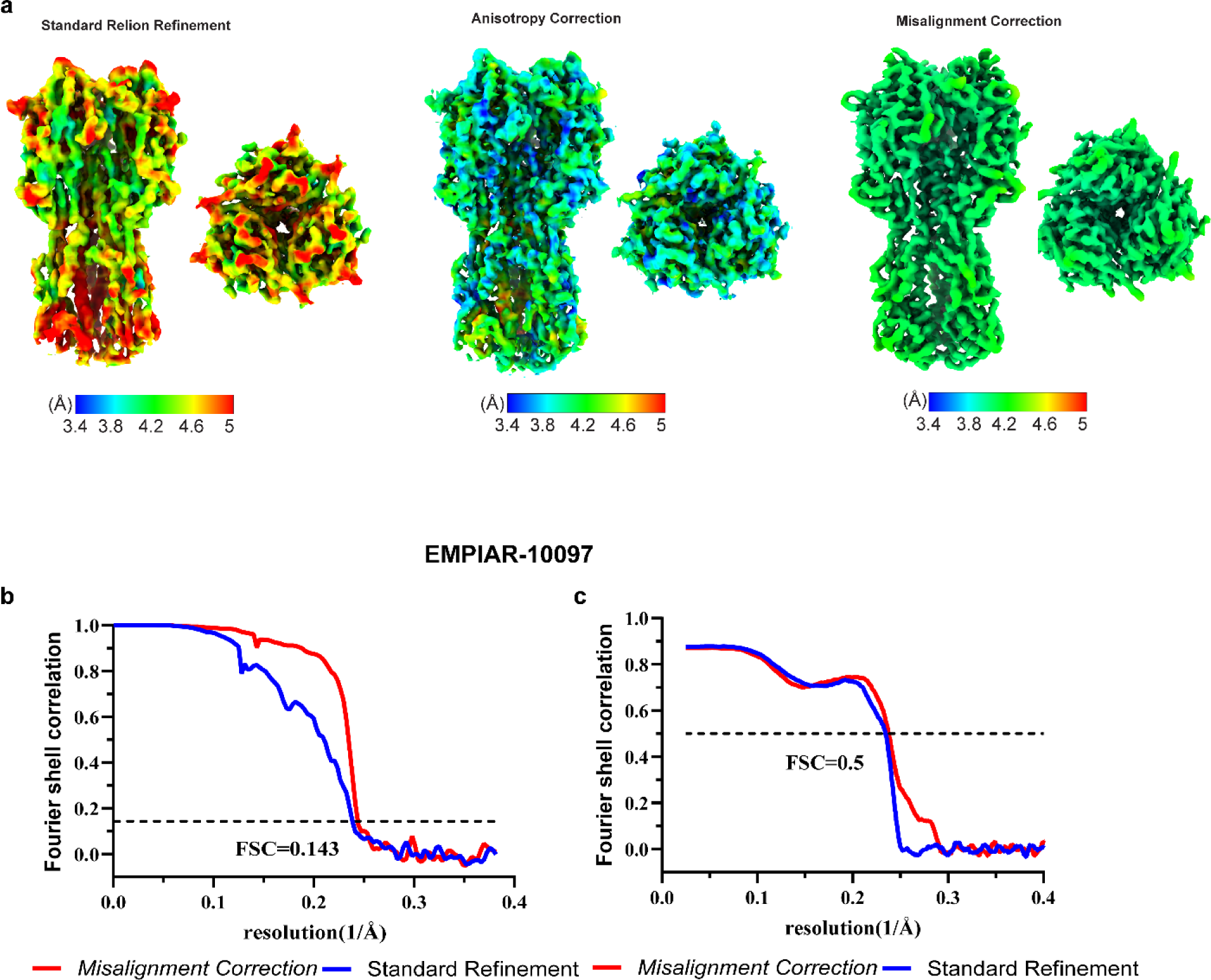
| spIsoNet *Anisotropy Correction and Misalignment Correction* on the HA trimer dataset collected with stage tilt. **a-b**, Local resolution map of HA trimer reconstructed by different methods: standard RELION Refinement, spIsoNet *Anisotropy Correction*, and spIsoNet *Misalignment Correction*. **b-c,** Half-map and model-to-map FSC curves of maps reconstructed with standard RELION Refinement and spIsoNet *Misalignment Correction*.

**Extended Data Fig. 4.**
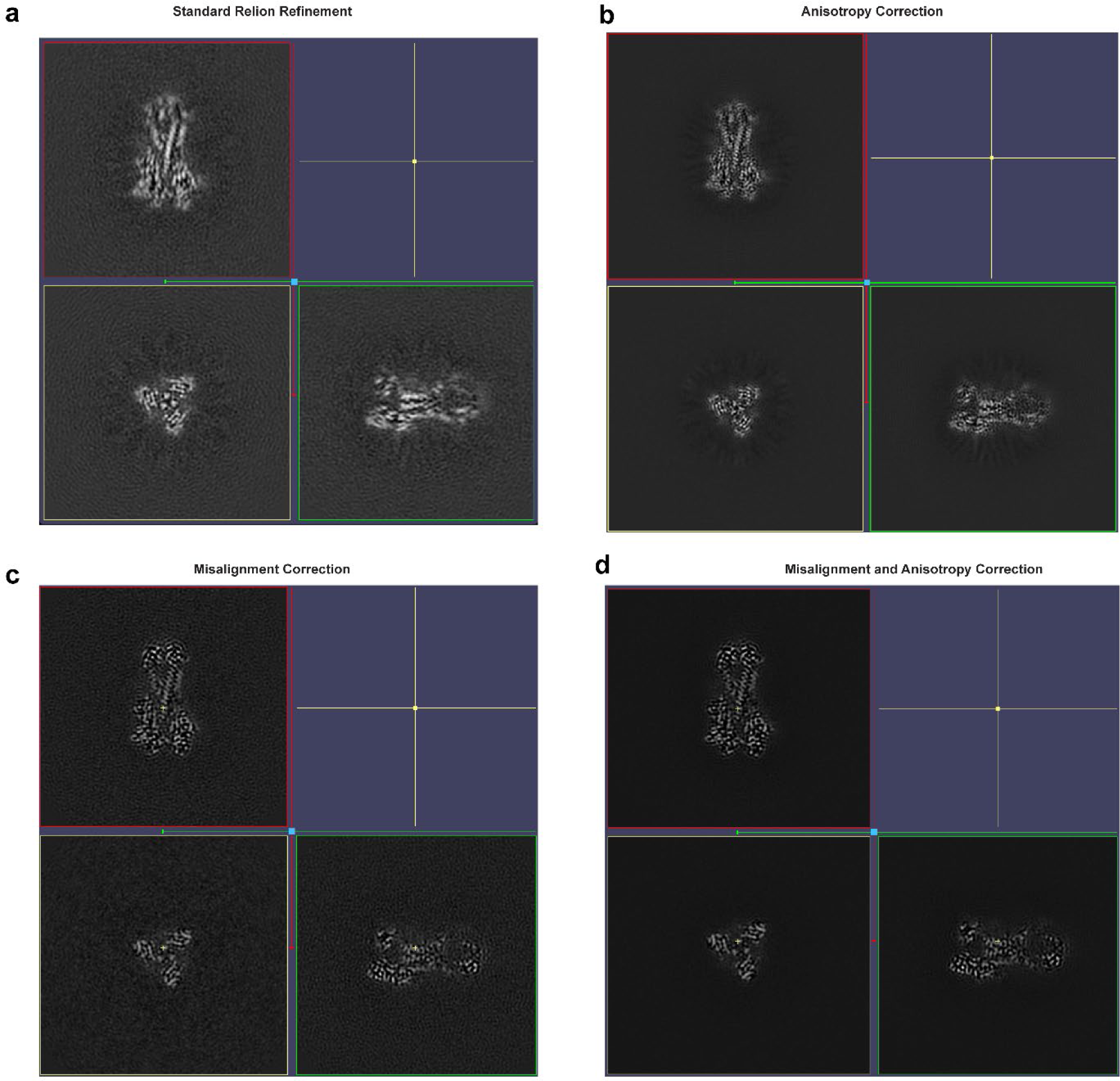
| Orthogonal map slices showing the effect of spIsoNet on the tilted cryoEM influenza hemagglutinin (HA) trimer dataset. **a-d**, The sections of orthogonal planes for the maps reconstructed by different refinement methods: standard RELION Refinement (**a**), *Anisotropy Correction* (**b**), *Misalignment Correction* (**c**), and the combination of *Misalignment and Anisotropy correction* (**d**).

**Extended Data Fig. 5.**
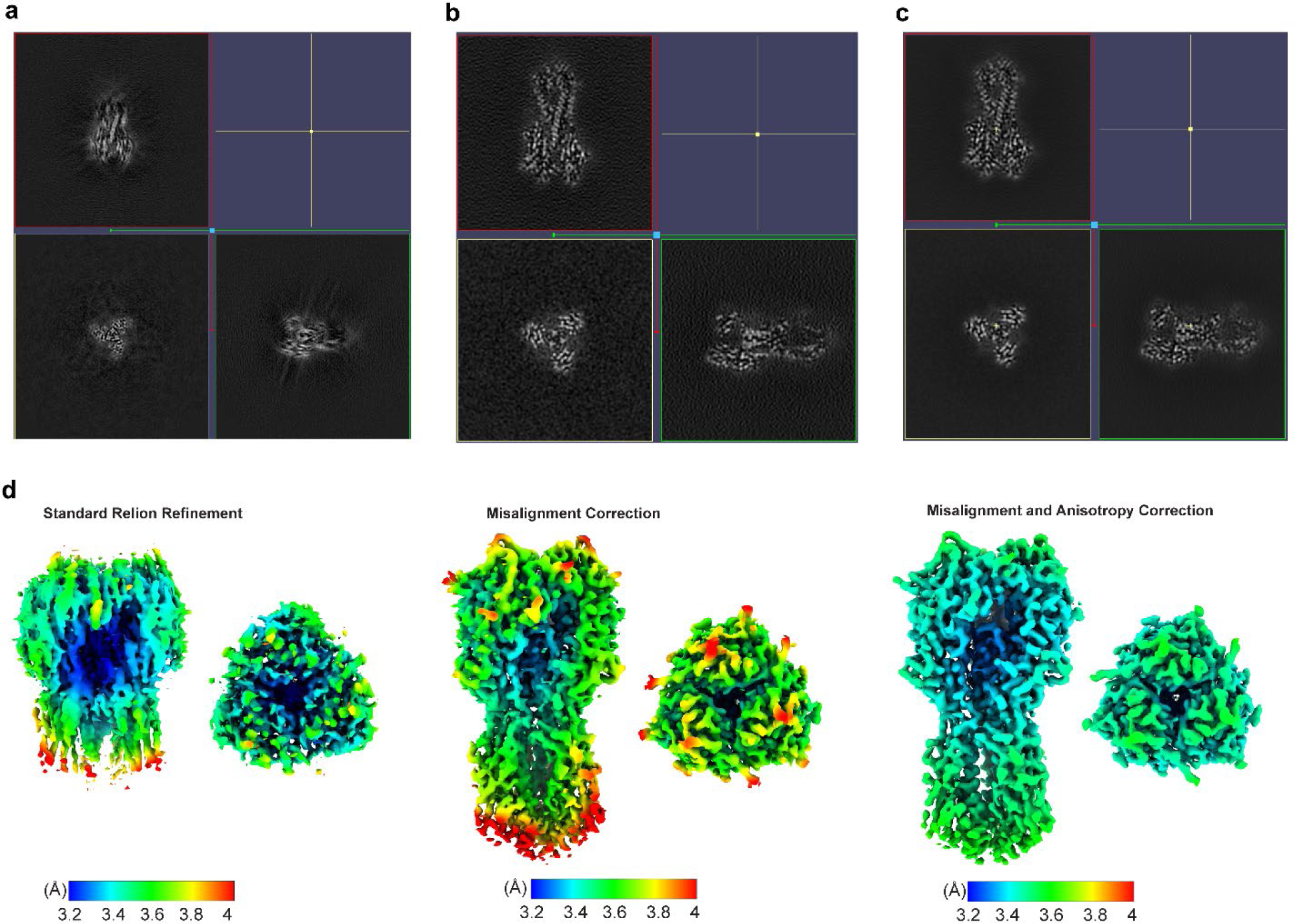
| The effect of spIsoNet on the untilted cryoEM influenza hemagglutinin (HA) trimer dataset. **a-c**, Orthogonal planes for the maps reconstructed by different methods: standard RELION Refinement (**a**), spIsoNet *Misalignment Correction* (**b**), and the combination of spIsoNet *Misalignment Correction* and *Anisotropy Correction* (**c**). **d**, Local resolution map of HA trimer reconstructed by different methods.

**Extended Data Fig. 6.**
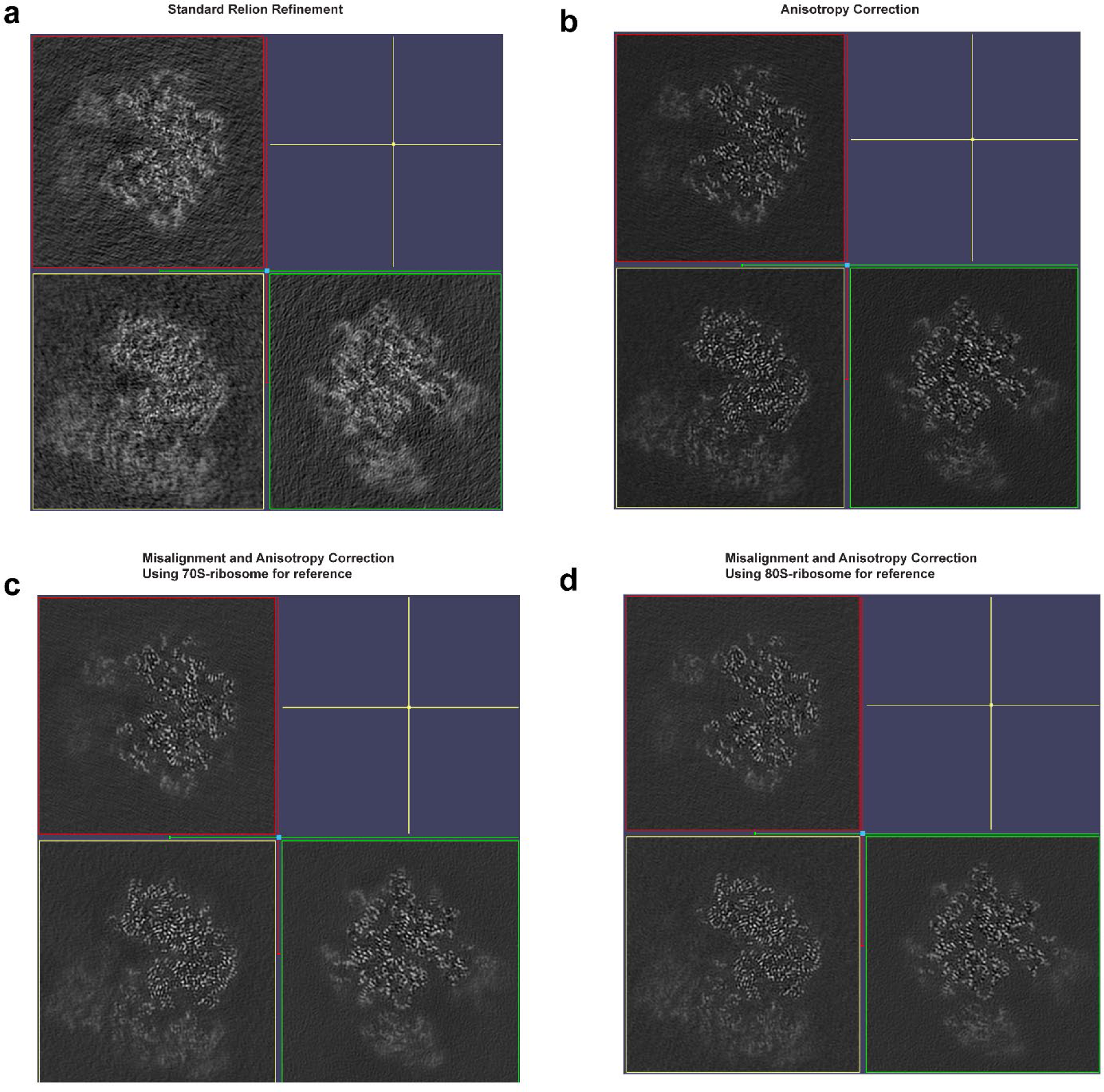
| Orthogonal slices showing effect of *Anisotropy and Misalignment Correction* on the experimental ribosome datasets. **a-d**, orthogonal sections of ribosome reconstructed by different methods: standard RELION refinement (**a**), spIsoNet *Anisotropy Correction* (**b**), spIsoNet *Misalignment Correction* using 70S ribosome as reference (**c**) and *Misalignment Correction* using 80S ribosome map as reference (**d**).

**Extended Data Fig. 7.**
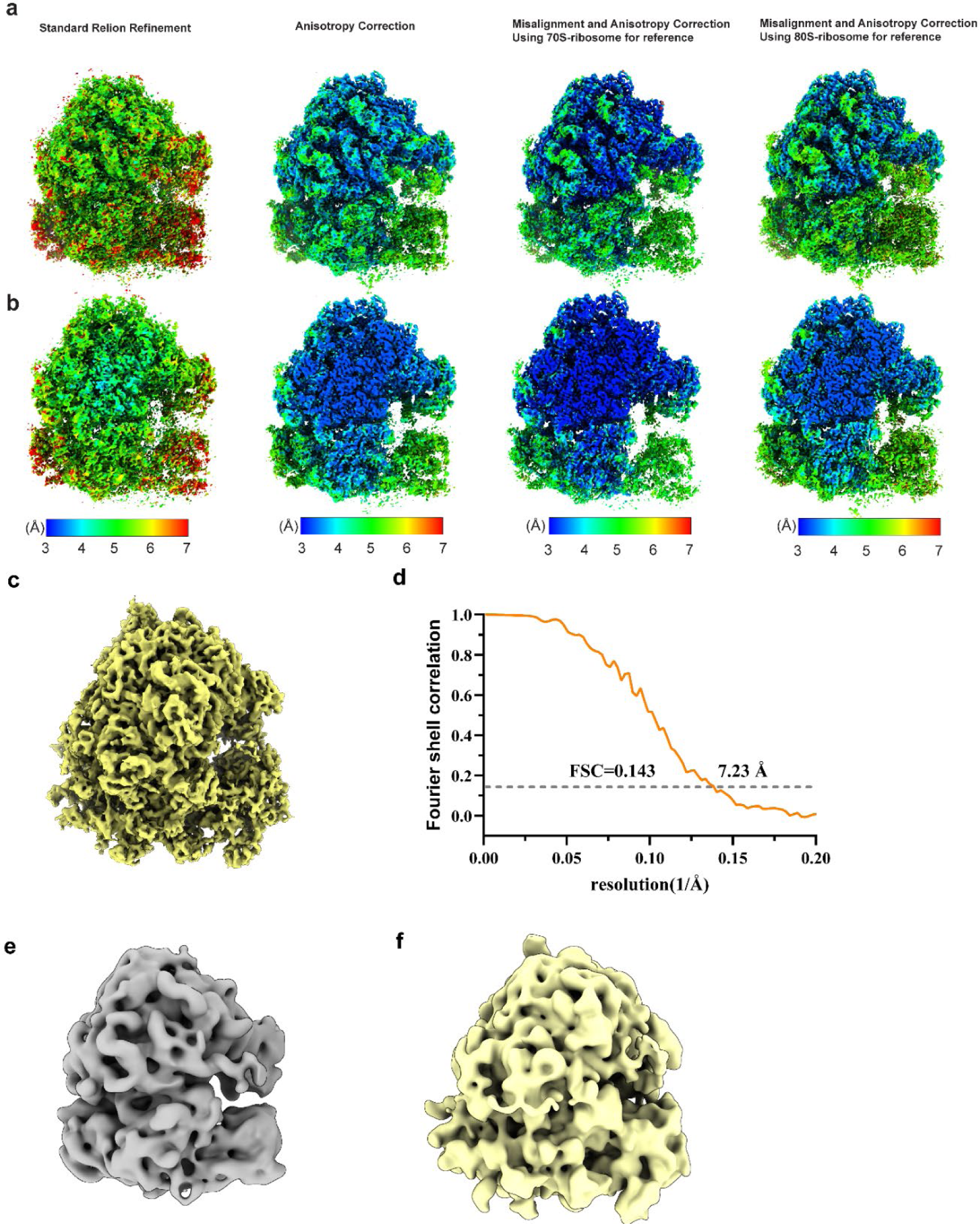
| spIsoNet *Anisotropy and Misalignment Correction* on the experimental ribosome datasets. **a-b**, Local resolution map of ribosome reconstructed by different methods: standard RELION refinement, spIsoNet *Anisotropy Correction*, spIsoNet *Misalignment Correction* and *Anisotropy correction* using 70S ribosome as reference, and *Misalignment Correction* and *Anisotropy correction* using 80S ribosome map as reference. **c-d,** Subtomogram average map of the 80S yeast ribosomes using EMPIAR 10045 with its FSC curves. **e-f,** The map of 70S (**e**) and 80S (**f**) low passed to 15Å and used as initial reference.

**Extended Data Fig. 8.**
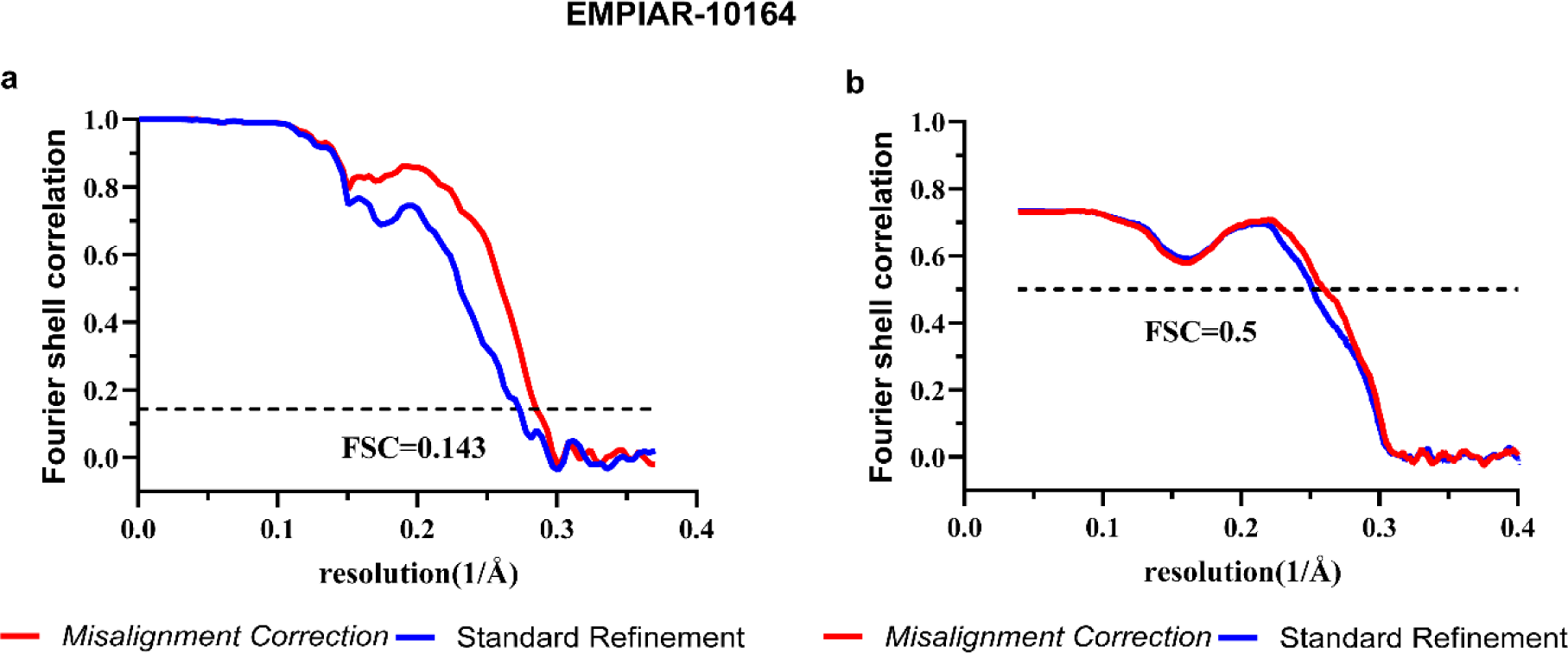
| Fourier Shell Correlation (FSC) plots for HIV-1 immature capsid datasets. **a,** The gold-standard FSC curves of the cryoEM HIV-1 immature capsid map with criterion of 0.143. Blue, RELION standard refinement; Red, spIsoNet *Misalignment Correction*. **b,** The Fourier shell correlation curves calculated between the cryoEM map and the refined atomic model with criterion of 0.5. Blue, RELION standard refinement; Red, spIsoNet *Misalignment Correction*.

